# The proto-oncogene DEK regulates neuronal excitability and tau accumulation in Alzheimer’s disease vulnerable neurons

**DOI:** 10.1101/2022.05.14.491965

**Authors:** Patricia Rodriguez-Rodriguez, Luis Enrique Arroyo-Garcia, Lechuan Li, Christina Tsagkogianni, Wei Wang, Isabella Salas-Allende, Zakary Plautz, Angel Cedazo-Minguez, Subhash Sinha, Olga Troyanskaya, Marc Flajolet, Vicky Yao, Jean-Pierre Roussarie

## Abstract

Neurons from layer II of the entorhinal cortex (ECII) are the first to accumulate tau protein aggregates and degenerate during prodromal Alzheimer’s disease. Here, we use a data-driven functional genomics approach to model ECII neurons *in silico* and identify the proto-oncogene DEK as a potential driver of tau pathology. By modulating DEK levels in EC neurons *in vitro* and *in vivo*, we first validate the accuracy and cell-type specificity of our network predictions. We then show that *Dek* silencing changes the inducibility of immediate early genes and alters neuron excitability, leading to dysregulation of neuronal plasticity genes. We further find that loss of function of DEK leads to tau accumulation in the soma of ECII neurons, reactivity of surrounding microglia, and eventually microglia-mediated neuron loss. This study validates a pathological gene discovery tool that opens new therapeutic avenues and sheds light on a novel pathway driving tau pathology in vulnerable neurons.

## INTRODUCTION

Early stages of neurodegenerative diseases are characterized by the aggregation of ubiquitous proteins in discrete populations of brain cells and degeneration of these cells. For most diseases this selective vulnerability pattern is unexplained yet could yield major insight into pathological mechanisms. This is the case for Alzheimer’s disease (AD), the world-leading cause of dementia. AD is defined by the appearance of two hallmark pathological lesions, amyloid plaques (extracellular aggregates of Aβ peptides) and neurofibrillary tangles (intracellular aggregates of hyperphosphorylated tau, or NFTs). While plaques are widespread in the neocortex and hippocampus, the appearance of NFTs follows a well-defined regional pattern that is commonly used to stage the disease and that starts in principal neurons from layer II of the entorhinal cortex (ECII) during prodromal AD (Braak & Braak, 1991). Very recently, positron emission tomography (PET) studies using tau tracers have provided a more complex picture, with AD clinical variants displaying different patterns of tau deposition. Nevertheless, AD patients with traditional amnestic presentation overwhelmingly fall into the classical entorhinal-centric NFT progression profile (Schöll et al., 2016; Vogel et al., 2021).

Neuroimaging studies in AD patients have shown that NFT is the AD feature that best correlates with regional cortical atrophy (Joie et al., 2020) and clinical symptoms of the disease (Bejanin et al., 2017; Ossenkoppele et al., 2016), suggesting the central role of tau in AD pathogenesis. The mechanisms responsible for the progression of NFT and neurodegeneration from ECII neurons to regions that are affected later in the disease remains to be fully understood, but the propagation of pathological tau along axonal routes has recently gained attention (Adams, Maass, Harrison, Baker, & Jagust, 2019; Vogel et al., 2020). Accordingly, misfolded tau seeds could spread across anatomically connected brain areas and convert normal tau from unaffected regions into neurotoxic aggregates. Before hippocampus and neocortex, the EC is the first region to develop such tau seeding activity (Kaufman, Del Tredici, Thomas, Braak, & Diamond, 2018). ECII neurons might thus form pathological tau *de novo* and could be a reservoir for tau spread to other regions. Understanding why ECII neurons accumulate tau during the earliest stages of AD could therefore reveal a major intervention point for disease modifying drugs.

The particularities of these cells have long remained elusive. Cell-type specific profiling technologies have made it possible to characterize their gene expression profiles, but how genes operate in the specific context of these cells and how they are related to disease cannot be readily parsed out by mere profiling. We recently used high quality molecular signatures of ECII neurons along with a large compendium of genomics data to build maps of functional gene interactions within the context of ECII neurons (Roussarie et al., 2020). Contrasting ECII neurons with neurons more resistant to AD, we found that pathways related to microtubule remodeling were particularly salient in ECII neurons. To investigate AD processes in the specific context of ECII neurons we used our NetWAS 2.0 (Network-Wide Association Study 2.0) algorithm, which leveraged our ECII functional map along with genome wide association study data for NFT formation (Beecham et al., 2014; Roussarie et al., 2020). NetWAS 2.0 re-prioritized genes based on their association with tau pathology within vulnerable neurons. This led to the identification of 4 functional modules potentially contributing to NFT formation in AD. Out of these 4 modules, one displayed higher connectivity in ECII neurons than in any other neuron type, thus it might represent a cellular process that is more central in ECII neurons than in other neurons. Furthermore, this module, enriched in genes involved in axonal plasticity, was affected by aging and amyloid pathology (Roussarie et al., 2020). While these results suggest that alterations in this module could underlie tau pathology in AD vulnerable neurons, details of its regulation, and its relevance for physiological and pathological mechanisms remain to be understood.

In the present work we build on this data-driven approach to identify a key driver of tau pathology within vulnerable neurons. We identify the DEK proto-oncogene as a hub gene of the AD vulnerability module and a close functional interaction partner of MAPT (tau protein gene). We first demonstrate that the effects of DEK on ECII neurons are accurately predicted by our *in silico* model. We further show that DEK, with previously unknown function in neurons, regulates immediate early gene (IEG) inducibility by modulating chromatin conformation. Downregulation of DEK consequently leads to alterations in neuronal excitability. This is accompanied by the accumulation and somatic redistribution of tau protein *in vitro*. Importantly, we show that the silencing of *Dek in vivo* leads to microglia reactivity and somatic buildup of tau in vulnerable neurons.

## RESULTS

### Identification of tau pathology drivers in ECII neurons using the NetWAS 2.0 systems biology approach

To determine potential drivers of tau pathology in ECII neurons, we honed in on the AD vulnerability module identified by the aforementioned NetWAS 2.0 approach (Roussarie et al., 2020). Specifically, we calculated the connectivity of every gene in the ECII functional network to genes in the AD vulnerability module (**Table 1**). Intuitively, high connectivity to genes in the AD vulnerability module suggests a strong functional relationship as a potential regulator. The gene with top connectivity to this module was *DEK*, suggesting that it might be central in the progression of tau pathology within vulnerable neurons. Furthermore, DEK was also predicted to be one of the closest functional interaction partners of MAPT, the gene that codes for tau (connectivity score 0.976, ranked 20 out of 23,950, data publicly available at alz.princeton.edu) (**Table 1**).

**Table 1.** Top 20 hits with highest connectivity scores to the NetWAS 2.0 vulnerability module and their respective associations with MAPT in the ECII neuron network.

|  | Connectivity to Netwas2.0 ECII neuron vulnerability module (Roussarie et al, 2020) | Average edge score to MAPT in ECII neuron network alz.princeton.edu |
| --- | --- | --- |
| DEK | 10.45 | 0.976 |
| TMX1 | 9.06 | 0.948 |
| PRKD3 | 8.94 | 0.763 |
| RECQL | 8.89 | 0.87 |
| MOB1A | 8.81 | 0.958 |
| EXOSC8 | 8.18 | 0.917 |
| SYPL1 | 8.18 | 0.996 |
| SMC4 | 8.17 | 0.844 |
| HAT1 | 8.14 | 0.954 |
| NDC1 | 8.13 | 0.693 |
| CCNA2 | 8.03 | 0.654 |
| ANP32E | 7.94 | 0.928 |
| HEATR1 | 7.94 | 0.922 |
| U2SURP | 7.92 | 0.961 |
| DIMT1 | 7.89 | 0.915 |
| NUP54 | 7.87 | 0.932 |
| SNAP23 | 7.84 | 0.886 |
| SRSF10 | 7.73 | 0.826 |
| PPP1CC | 7.72 | 0.866 |
| NUP107 | 7.68 | 0.851 |

*DEK* codes for the proto-oncogene DEK, a nuclear phosphoprotein that was identified for the first time in patients with acute myeloid leukemia (Von Lindern et al., 1992) and is frequently upregulated in solid tumors such as melanoma and breast cancer (Carro et al., 2006). It has also been associated with several autoimmune diseases, including lupus erythematosus and juvenile arthritis (Dong et al., 2000). DEK is ubiquitously expressed in the central nervous system **(Figure S1)**, although its function there remains largely unknown. Its importance in a wide variety of diseases, combined with our computational analyses that implicate it in processes underlying ECII vulnerability, motivated us to experimentally characterize DEK function in ECII neurons, and its role in pathological processes leading to AD.

### *Dek* silencing in EC neurons leads to alterations in AD associated pathways

To study DEK in the context of vulnerable neurons, we modulated *Dek* expression in mouse EC neuron primary cultures by transducing them with adeno-associated viruses (AAVs) carrying either a cDNA coding for *Dek*, a silencing small hairpin RNA (shRNA) directed against *Dek*, or their respective controls. Transduction with these viruses led to a very significant overexpression and silencing of *Dek*, respectively (overexpression logFC = 1.79, FDR= 4.33E-80; silencing logFC= −2.18, FDR = 1.55E-58) (**Table S1**). RNA-seq followed by differential gene expression (DGE) analysis 4 days after transduction showed that *Dek* silencing has larger effects on the transcriptional landscape of the cells than *Dek* overexpression (**Figure 1a, 1b, 1d and 1e**). Pathway analysis (Ingenuity pathway analysis, DEGs FDR< 0.05) revealed that some of the most significantly downregulated pathways in *Dek*-silenced neurons are associated with microtubule dynamics (p-value = 2.94E-62), development of neurons (p-value = 1.41E-49), and axonogenesis (p-value = 1.50E-19). The synaptic longterm potentiation (LTP) signaling pathway was also significantly decreased (p-value = 5.09E-21), suggesting alterations in synaptic transmission (**Figure 1c**). Considering the modest number of significantly modified genes upon *Dek* overexpression by adjusted p value, we decided to perform pathway analysis on genes with nominally significant p-values (p-value <0.05). These results showed that similar pathways, including LTP (p-value = 7.61E-5), microtubule dynamics (p-value = 2.71E-11) and development of neurons (p-value = 3.07E-8), were also consistently altered in *Dek*-overexpressing neurons (**Figure 1f**) supporting the notion that DEK could play a role as a modulator of synaptic transmission and plasticity in EC neurons.

**Figure 1.**
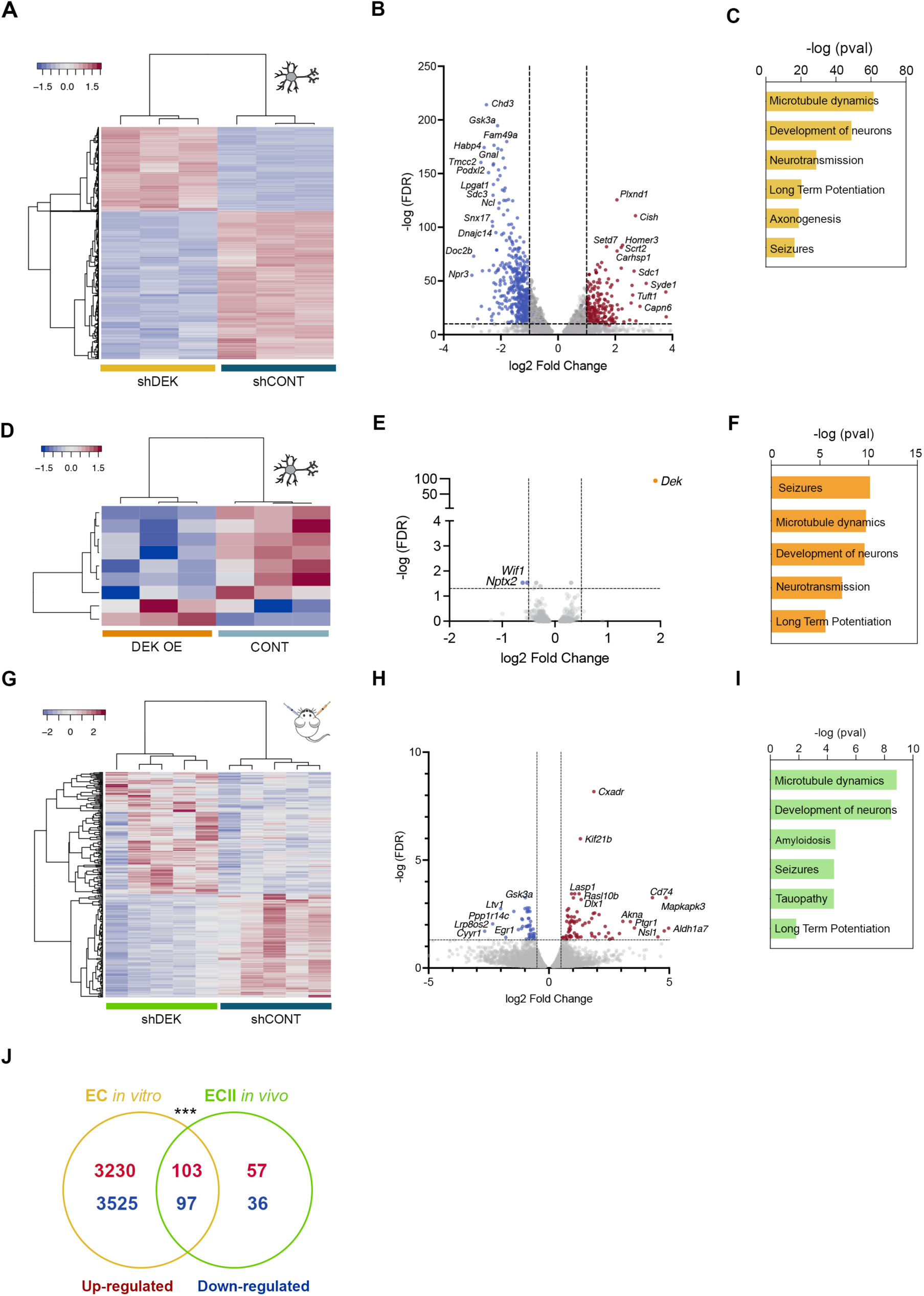
Gene expression changes in EC neurons triggered by modulation of DEK in vitro and in vivo. **A)** Heatmap of the DEGs between control and Dek-silenced primary neurons. **B)** Volcano plot of DEGs between control and Dek-silenced primary neurons. **C)** Pathways that are altered in Dek-silenced primary neurons compared to control. **D)** Heatmap of the DEGs between control and Dek-overexpressing primary neurons. **E)** Volcano plot of DEGs between control and Dek-overexpressing primary neurons. **F)** Pathways that are altered in Dek-over-expressing primary neurons compared to control. **G)** Heatmap of the DEGs between control and Dek-silenced ECII neurons in vivo. **H)** Volcano plot of DEGs between control and Dek-silenced ECII neurons in vivo. **I)** Pathways that are altered in Dek-silenced ECII neurons in vivo compared to control. **J)** Venn diagram that shows a significant overlap of up and downregulated DEGs between Dek-silenced neurons in vitro and in vivo. Fisher’s exact test pval < 0.0001.

To study DEK function *in vivo*, we silenced *Dek* in ECII neurons of ECII-bacTRAP mice previously generated in our laboratory. ECII-bacTRAP mice express a fusion protein between EGFP and the ribosomal protein L10a in ECII neurons specifically. Actively translated mRNA from these neurons can be isolated by immunoprecipitating enhance green fluorescent protein (EGFP) from EC lysates and profiled by RNA-seq (Heiman, Kulicke, Fenster, Greengard, & Heintz, 2014; Roussarie et al., 2020). Perturbing *Dek* expression in the EC of these mice allows to measure the effects of DEK on gene expression regulation specifically in ECII neurons. BacTRAP-RNAseq analysis one week after AAV injection confirmed the successful silencing of *Dek* in ECII neurons (logFC = −0.69, p-value = 9.9E-04) **(Figure 1g, 1h; Table S1)**. In agreement with the *in vitro* data, pathway analysis (Ingenuity pathway analysis, DEGs FDR< 0.05) revealed again alterations in cellular functions related to microtubule dynamics (p-value = 1.40E-09), development of neurons (p-value = 3.38E-9) and LTP (p-value = 0.015) in *Dek* silenced ECII neurons compared to the control (**Figure 1i**). We found a significant overlap between DEK regulated genes *in vitro* and *in vivo* (**Figure 1j**).

Analysis of DEK-regulated genes in EC neurons allowed us to test the accuracy of the predictions of our ECII network (the *in silico* network described above). Genes predicted to be highly functionally connected to DEK in ECII neurons were indeed more affected by *Dek* silencing in EC neurons *in vitro* than expected by chance (p-value < 2.2E-16, one-sided Wilcoxon rank sum test of DEGs FDR < 0.05, **Figure S2**). The ECII functional network was also highly predictive for genes differentially expressed after *Dek* silencing *in vivo* (p-value = 8.09E-15, one-sided Wilcoxon rank sum test of DEGs FDR < 0.05, **Figure 2**). The ECII functional network is thus a powerful discovery tool for the unbiased identification of regulatory processes. Furthermore, we found that genes from the AD vulnerability module were more differentially expressed after *Dek* silencing than expected by chance (p-value = 3.3E-09, Fisher’s exact test), thereby validating DEK as a significant regulator of the AD vulnerability gene module. Altogether, these data demonstrate experimentally that, in the context of ECII neurons, DEK indeed regulates genes predicted by NetWAS 2.0 to be associated with ECII vulnerability to AD.

**Figure 2.**
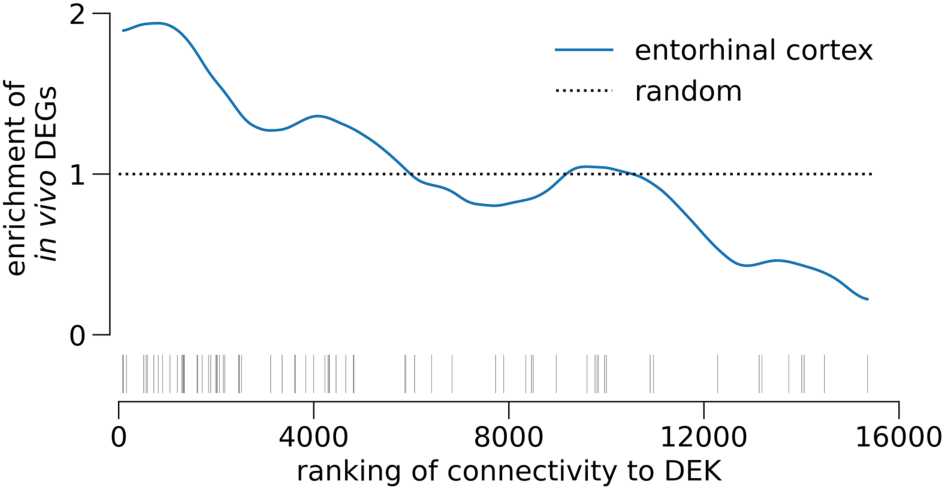
Significant enrichment of genes changing in vivo from Dek-silencing in in silico ECII network predictions. Enrichment analysis of DEGs (FDR<0.05, denoted in the rug plot, bottom) between control and Dek-silenced ECII neurons in vivo demonstrates strong enrichment within genes ranked by probability of functional interaction with DEK in the entorhinal cortex network (blue). Dashed line represents expected enrichment if given random predictions.

### *Dek* silencing in EC neurons alters the expression of IEGs

DEK has been previously identified as a chromatin-binding protein that plays a variety of roles in the regulation of chromatin structure and gene expression (Kappes, Scholten, Richter, Gruss, & Waldmann, 2004; Sandén et al., 2014). To explore whether the effect of DEK on neuronal function is accompanied by alterations in chromatin accessibility, we performed ATAC-seq (assay for transposase-accessible chromatin using sequencing) in primary cultures of EC neurons, 4 days after *Dek* silencing. Our results showed differences in chromatin accessibility at the promoter regions of a total of 20 genes (**Table S2**). These alterations were accompanied by differential expression of 13 of these genes, determined by our previous *in vitro* RNAseq analysis (**Figure 3a**). Interestingly, as shown in **Figure 3a**, the IEG *Egr1* was highly significantly downregulated (FDR = 4.12E-48), with the largest fold change (log FC = −1.85) among these genes. Furthermore, *Egr1* was the only one that was also differentially expressed (downregulated, log FC = −0.76) in *Dek*-silenced ECII neurons *in vivo* (FDR = 0.018), as shown in our bacTRAP-RNAseq data (**Table S1**). Surprisingly though, the concomitant downregulation of *Egr1* and increase in chromatin accessibility at its proximal enhancer and promoter regions (FDR = 0.0417) (**Figure 3b**) was counterintuitive and led us to investigate further the regulation of IEGs in *Dek*-silenced neurons.

**Figure 3.**
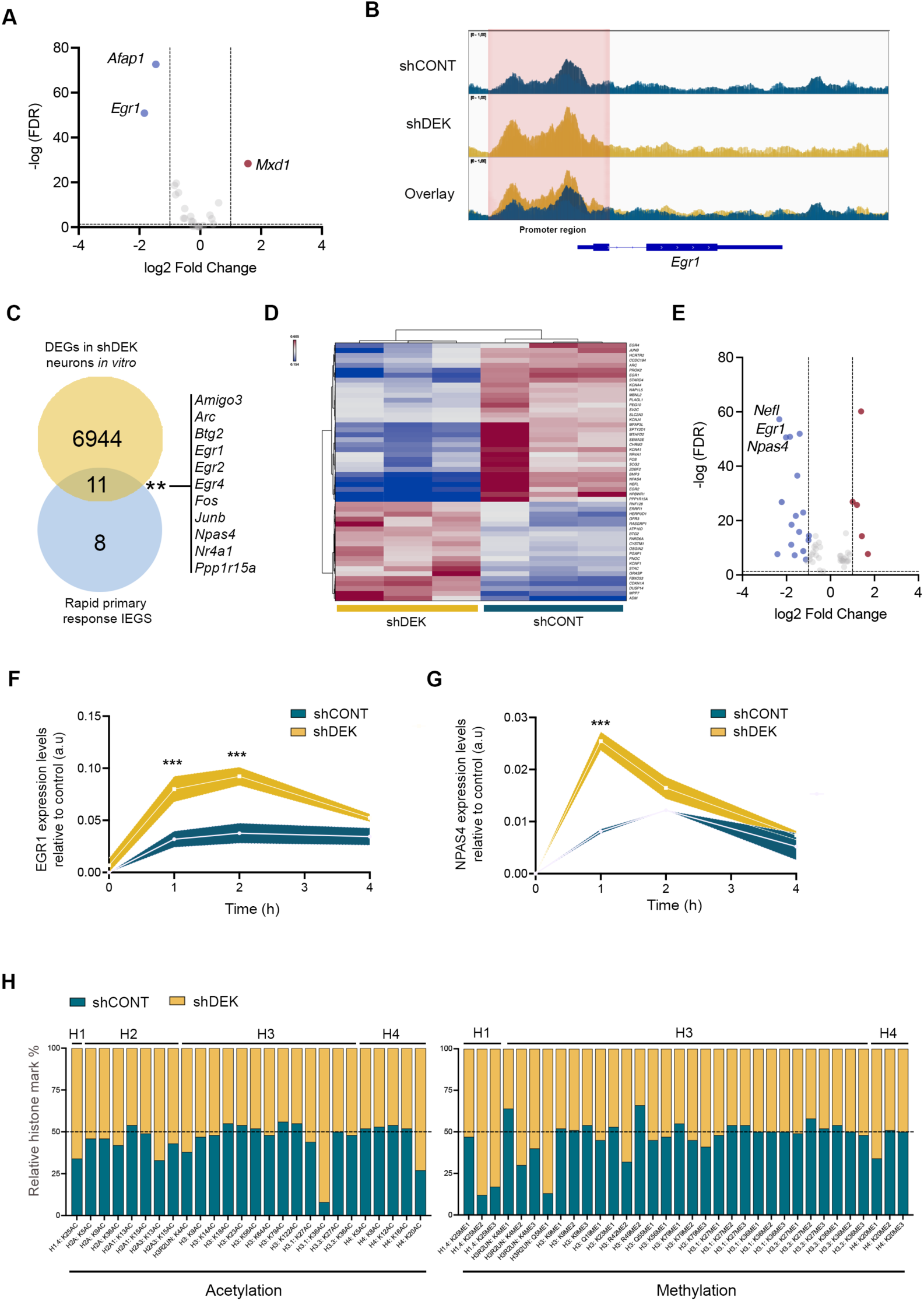
Effect of Dek silencing on chromatin accessibility, immediate early gene expression and histone marks. **A)** Volcano plot of the DEGs in vitro that also showed alterations in chromatin accessibility at their loci. **B)** ATACseq tracks of Dek-silenced (yellow), control neurons (blue) and their overlay. **C)** Venn diagram of the overlap between rapid primary response IEGs and DEGs in Dek-silenced neurons in vitro. Fisher’s exact test *pval <0.05. **D)** Heatmap of all the IEGs that are differentially expressed in Dek-silenced neurons compared to control. **E)** Volcano plot of all the IEGs that are differentially expressed in Dek-silenced neurons compared to control. **F)** Expression levels of Egr1 after 1, 2, 3 and 4 hours of induction with 50 mM KCl in control and Dek-silenced neurons. 2way ANOVA Sidak’s multiple comparisons test *** pval<0.005. **G)** Expression levels of Npas4 after 1, 2, 3 and 4 hours of induction with KCl in control and Dek-silenced neurons. 2way ANOVA Sidak’s multiple comparisons test *** pval<0.005. **H)** Relative abundance of acetylated and methylated histone marks in control and Dek-silenced neurons.

IEGs, like Egr1, are the first genes induced upon neuronal activation and are essential modulators of long-term neuronal plasticity and learning (Bozon, Davis, & Laroche, 2003; Jones et al., 2001; Yap & Greenberg, 2018). Among IEGs, a first wave of genes is transcribed immediately upon neuronal activation while a second wave is transcribed after the activation of these primary genes. We found a significant overlap between genes differentially expressed after *Dek* silencing and a published set of rapid primary response IEGs (Tyssowski et al., 2018) **(Figure 3c)**. Delayed primary response genes, were also significantly affected, although not as consistently as the rapid primary response genes **(Figure 3d)**. Out of all IEGs, the top downregulated IEGs were the primary response genes *Egr1* (FDR = 4.12E-48) and *Npas4* (FDR = 9.71E-50) (**Figure 3e**). To test the direct effects of DEK on IEG induction, we used 50 mM KCl to cause neuronal depolarization and calcium influx in a manner that is largely independent from synaptic transmission.

We then monitored the expression levels of *Egr1* and *Npas4* over time using quantitative PCR. As expected, KCl caused a large transient increase in Egr1 and Npas4 expression in control neurons. Strikingly though, KCl led to a maximum 2.5- and 3-fold higher induction of *Egr1* and *Npas4* expression respectively in EC neurons where *Dek* was silenced compared to control (**Figure 3f and 3g**). The larger induction of *Egr1* is in agreement with higher chromatin accessibility at the proximal enhancer and promoter regions of the gene and suggests that the downregulation of *Egr1* at baseline is due to secondary alterations in neuronal activity. On the other hand, the ATAC-seq did not show significant effects of *Dek* silencing on chromatin accessibility at the *Npas4* locus, indicating additional mechanisms that regulate IEG expression by DEK.

Epigenetic modifications at the cis regulatory elements of IEGs are essential to regulate their expression (L.-F. Chen et al., 2019; Malik et al., 2014) and a number of studies showed a role for DEK as a histone chaperone and modulator of histone modifications (Hollenbach, McPherson, Mientjes, Iyengar, & Grosveld, 2002; Ko et al., 2006). In particular, DEK was suggested to mediate the assembly of H3.3 nucleosomes (Ivanauskiene et al., 2014; Sawatsubashi et al., 2010). To test if *Dek* silencing would alter the location of H3.3 nucleosomes in the genome, we used the CUT&Tag technique (cleavage under targets and tagmentation), whereby an antibody against H3.3 is coupled to a tagmentation enzyme to introduce DNA adapters at the binding locations. DNA sequencing then allows mapping of H3.3 nucleosomes based on the genomic sequences adjoining the adapters. The CUT&Tag assay in control and *Dek*-silenced EC neurons 4 days after transduction revealed only one gene that presented significant differences in H3.3 accumulation, *Lrrc4c*. We could not find any difference in H3.3 deposition at the *Egr1* or *Npas4* loci. Neuronal activity induced acetylation of histone H3 at lysine 27 (H3K27ac) at the enhancers of IEGs regulates their transcription (L.-F. Chen et al., 2019; Xiaohui Li et al., 2021; Malik et al., 2014), and a previous study showed that DEK chromatin binding pattern overlaps with H3K27ac (Sandén et al., 2014). To determine if *Dek*-silencing was modulating H3K27ac deposition at the *Egr1* and *Npas4* loci we performed another CUT&Tag assay. Our results revealed differences in H3K27ac levels at a total of 779 genome locations in *Dek*-silenced neurons compared to control. However, we did not observe differences in its signal around *Egr1* or *Npas4*.

To test in an unbiased manner if other specific histone marks could be modulated by DEK, we conducted a mass spectrometry-based screen of histone posttranslational modifications in control and *Dek*-silenced EC neurons. The assay revealed differences in total levels of H4K20ac, H1.4K25me2, H1.4K25me3 and H3Q5me1 (2.7, 7.3, 4.9, 6.7-fold increase respectively) and a striking more than 10-fold increase in H3.1K36ac levels in *Dek*-silenced neurons compared to control (**Figure 3h**). To test if the increased inducibility of IEGs after *Dek* silencing could be mediated by H3.1K36ac we performed another a CUT&Tag assay. Unfortunately, currently available antibodies cannot discriminate between H3.1K36ac and H3.3K36ac (that shows 6-times higher levels than H3.1K36ac in our neurons and is not changed by *Dek*-silencing), making it challenging to see H3.1K36ac specific changes. We found no differences in H3K36ac deposition around the *Egr1* or *Npas4* loci (**Table S2**) in *Dek*-silenced neurons compared to control. Mapping genomic locations for the other above-mentioned histone marks affected by DEK is also limited by the lack of availability of reagents.

### *Dek* silencing in EC neurons *in vitro* leads to changes on intrinsic excitability

The modulation of IEGs by DEK could have profound functional consequences on *Dek*-silenced neurons. To test this hypothesis, we performed patch-clamp recordings in *Dek*-silenced EC neurons in primary culture 4 days after transduction. Evaluation of the resting membrane potential of *Dek*-silenced neurons showed that they were significantly depolarized compared to the control group (**Figure 4a and 4b)**. Concurrently, the rheobase, or minimal current required to elicit an action potential, was significantly lower, suggesting increased excitability (**Figure 4c**). Furthermore, we found decreased membrane input resistance, and increased firing threshold (**Figure 4d and 4e**) that confirm that the neuronal intrinsic properties are altered by *Dek* silencing (**Table S3**). These results suggest a functional change in Na^+^ currents. Indeed, our RNAseq *in vitro* data showed a broad dysregulation of sodium channel expression, with highly significant downregulation of pore-forming subunits of voltage-gated sodium channels Nav1.2 (*Scn2a)* Nav1.6 (*Scn8a*) and Nav1.1 (*Scn1a*), and upregulation of accessory subunits of Nav channels *Scn1b* and *Scn3b* (**Figure 4f-k**). While the downregulation of voltage gated sodium channels could seem at odds with the increased excitability observed upon *Dek* silencing, a recent study also found increased excitability after invalidation of Nav1.2 (Spratt et al., 2021).

**Figure 4.**
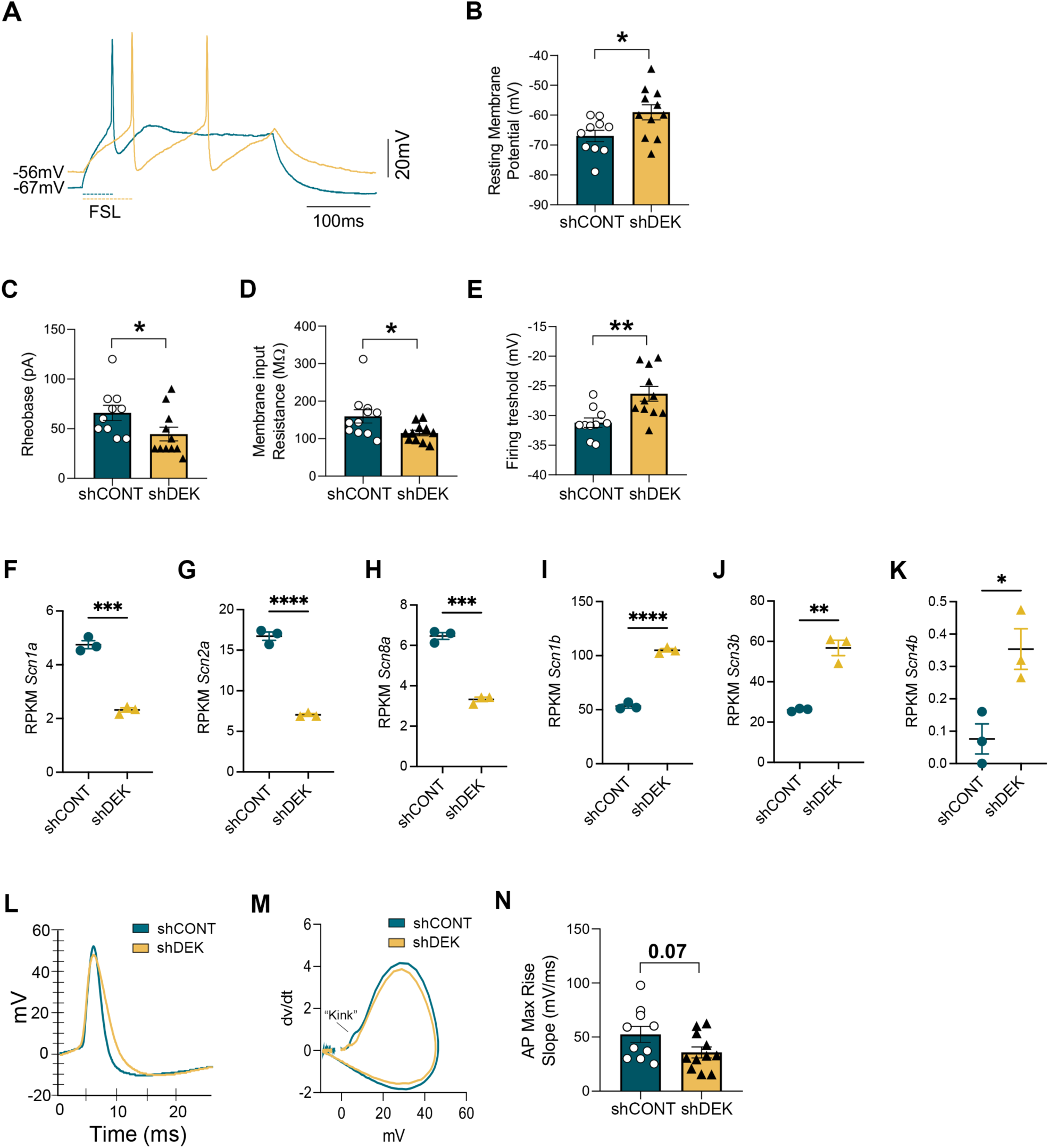
Electrophysiological properties of Dek-silenced EC primary neurons. **A)** Representative traces of first spike and resting membrane potential from: control (blue) and Dek-silenced neurons (yellow). **B**) Bar graph of the resting membrane potential in control vs Dek-silenced neurons. Unpaired t-test *pval<0.05. **C**) Bar graph of the rheobase in control vs Dek-silenced neurons. Mann-Whitney test *pval<0.05. **D)** Bar graph of the membrane input resistance in control vs Dek-silenced neurons. Mann-Whitney test *pval<0.05. **E)** Bar graph of the firing threshold in control vs Dek-silenced neurons. Unpaired t-test *pval=0.08. **F-K)** Expression levels of Scn1a, Scn2a, Scn8a, Scn1b, Scn3b and Scn4b in control vs Dek-silenced neurons. **L)** Representative action potential (AP) waveform in control and Dek-silenced neurons. **M)** Representative phase plot in control and Dek-silenced neurons. **N)** Bar graph of the AP maximum rise slope in control vs Dek-silenced neurons. Unpaired t-test *pval=0.07.

The conductance of voltage-gated sodium channels can be evaluated by determining the kinetics of the action potential (AP) initiation. We thus analyzed the AP waveform and phase plot (Platkiewicz & Brette, 2010; Prestigio et al., 2019). We did not find significant alterations in the AP waveform (**Figure 4l**). However, we observed a lack of the first spike initiation, or “kink” (**Figure 4m**). This result coincides with a decrease in the AP max rise slope (**Figure 4n**) and suggest a change in the density of voltage-gated sodium channels localized at the neuronal axon initial segment (Prestigio et al., 2019).

### *Dek* silencing leads to tau accumulation and ECII neuron loss

A strong body of evidence suggests that tau accumulation and neuronal excitation are closely linked. Pathological tau can affect the organization of the axon initial segment and disrupt excitability homeostasis (Hatch, Wei, Xia, & Götz, 2017; Sohn et al., 2019, 2016). Conversely, sustained neuronal excitation can lead to increased tau accumulation (Siddhartha et al., 2018; Wu et al., 2016). Considering this evidence, and since our data-driven analysis initially predicted that DEK could be central to tau pathology, we then asked if the modulation of neuronal excitability by DEK could be associated with alterations of the tau protein. We first tested the effect of DEK on tau in mouse primary EC neurons. We measured total tau protein levels and its phosphorylation by western blot at different time points following *Dek* silencing. As shown in **Figure 5a**, *Dek* silencing led to increased tau levels at day 4 after transduction. This was not accompanied by increased tau phosphorylation at Ser202/Thr205 (as measured by the AT8 antibody), generally associated with early AD pathology (Luna-Muñoz, Chávez-Macías, García-Sierra, & Mena, 2007; Su, Cummings, & Cotman, 1994). We measured tau mRNA and found that higher tau protein levels were not paralleled by tau mRNA levels, suggesting that DEK leads to tau accumulation by modulating its translation, release, or degradation (**Figure 5b**). Lastly, tau abnormal redistribution to the somatodendritic compartment is an early event in AD pathology (C. Li & Götz, 2017; Xiaoyu Li et al., 2011). Using immunofluorescence at 4 days post-transduction, we confirmed the increase in total tau levels observed by western blot and showed that *Dek* silencing leads to the relocation of tau protein in the neuronal soma (**Figure 5c**).

**Figure 5.**
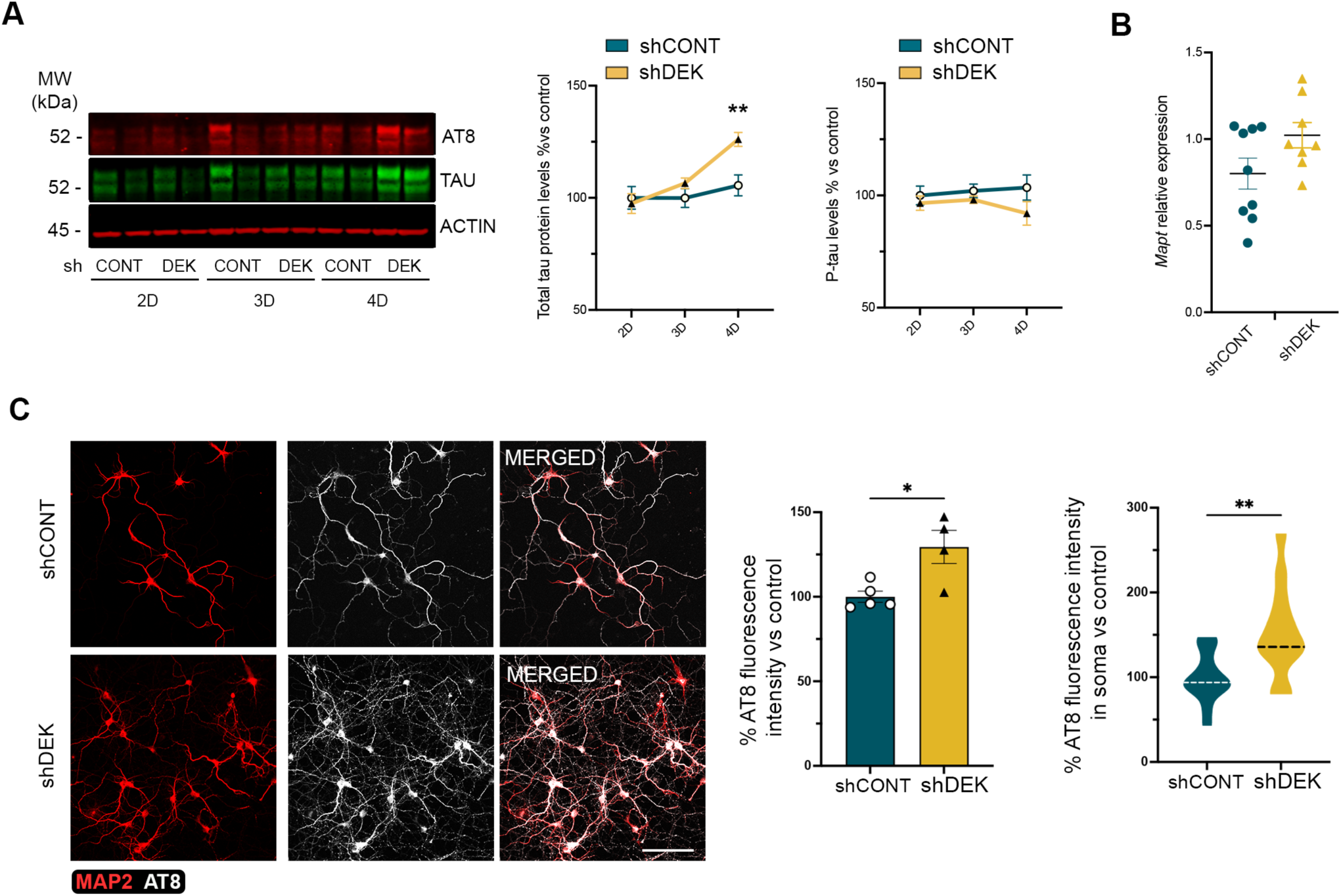
Effect of Dek silencing in EC neurons in vitro. **A)** Western blot analysis of the levels of total tau (green) and phospho-tau Ser202/Thr205 (AT8, red) in EC neurons in primary culture at 2-, 3- and 4-days post-transduction. The graphs show protein levels in % relative to 2 days post-transduction control neurons. Band intensity was normalized with actin for total tau measurements and with total tau for phospho-tau measurements. 2way ANOVA Sidak’s multiple comparisons test **pval<0.01. **B)** RT-qPCR analysis of Mapt expression levels in control and Dek-silenced neurons 4 days after transduction. **C)** Confocal microscopy images of immunofluorescence staining of MAP2 (red) and AT8 (greyscale) in control and Dek-silenced neurons at 4 days post-transduction. Scale bar 100 µm. The graphs show AT8 fluorescence intensity levels in % relative to control-treated neurons. Un-paired t-test **pval<0.01, *pval<0.05.

To determine the effects of *Dek* silencing *in vivo*, we performed stereotaxic injections of a control- and a *Dek* silencing-AAV in opposite entorhinal cortex hemispheres of human-tau transgenic mice (hMAPT mice). This mouse model is knocked-out for mouse tau while expressing the full length human MAPT gene, and can generate all human tau isoforms, including 4R and 3R tau (Andorfer et al., 2003). Immunofluorescence labeling of ECII neurons with Reelin at 4 days, 1 week and 2 weeks after injection revealed a highly significant loss of these neurons after 2 weeks (**Figure 6a**). Iba1 immunofluorescence staining of microglia showed a significant increase in reactive microglia specifically in the layer II of EC at 2 weeks post-injection (**Figure 6a**). At 1-week post-injection (before neuron loss), we found no differences in tau protein levels in ECII by immunofluorescence against phospho-tau (Thr231) (**Figure S3a**). As we observed an accumulation of reactive microglia around transduced neuron cell bodies **(Figure S3b)**, we hypothesized that microglia might be an important contributor to the neuron loss observed at 2 weeks post-injection. Indeed, RT-qPCR on EC lysates showed a significant increase in expression of the microglia markers *Cx3cr1, Itgam* (that codes for CD11b) and complement C*1qa* (associated with synaptic pruning) (Lui et al., 2016; Vasek et al., 2016). We also found an increase in disease-associated microglia genes *Tyrobp* and *Spp1* (Mathys et al., 2019; Olah et al., 2020) *a*s well as in *Ccl3*, upregulated in microglia in response to neuronal activation (Badimon et al., 2020),. We did not observe significant differences in expression of *Ccl24* or *P2ry12* (also upregulated in microglia in response to neuronal activation) (Badimon et al., 2020) **(Figure S4)**.

**Figure 6.**
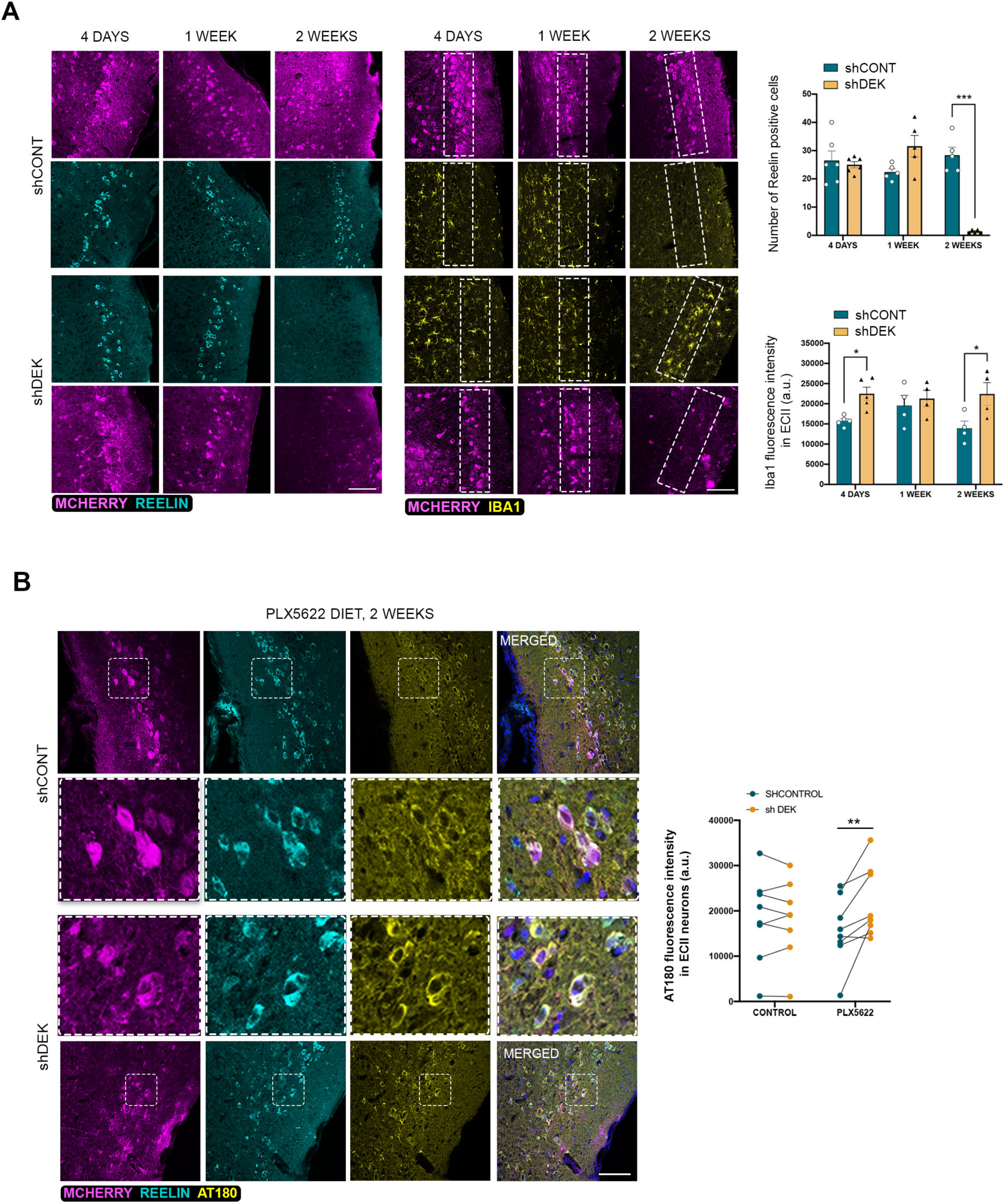
Effect of Dek silencing in ECII neurons of hMAPT mouse. **A)** Confocal microscopy images of immunofluorescence staining of transduced neurons (mCherry, magenta), ECII neurons (Reelin, cyan) and microglia (Iba1, yellow) at 4 days, 1 week and 2 weeks post-transduction of hMAPT mice. Scale bar 100 µm. The graphs show the quantifications of the number of reelin positive cells and of Iba1 fluorescence intensity in layer II of the mouse entorhinal cortex in the control and Dek-silenced hemi-spheres. 2way ANOVA Sidak’s multiple comparisons test ***pval<0.005, *pval<0.05. **B)** Confocal microscopy images of immuno-fluorescence staining of transduced neurons (mCherry, magenta), ECII neurons (Reelin, cyan) and phospho-tau Thr231 (AT180, yellow) at 2 weeks post-transduction of control or shDek-carrying AAVs in PLX5622 diet fed hMAPT mice. Scale bar 100 µm. The graph shows the quantification of AT180 fluorescence intensity, paired by the control and the Dek-silenced opposite hemisphere for each hMAPT mouse. 2way ANOVA Sidak’s multiple comparisons test **pval<0.01.

To test the possibility that longer-term silencing of DEK could lead to an accumulation of tau like that seen *in vitro*, we attempted to eliminate microglial cells, to increase ECII neuron survival. For that purpose, we fed mice with chow containing PLX5622. PLX5622 is an antagonist of colony-stimulating factor 1 receptor (CSF1R), that is essential for microglia survival. Treatment of mice with this compound leads to microglia depletion after 5 days (Spangenberg et al., 2019). At day 5 we injected control or sh*Dek* AAVs in the EC of opposite hemispheres and we analyzed ECII neuron survival and tau accumulation at 2 weeks post-injection **(Figure S5a**). As shown in **Figure S5b and S5c**, PLX5622 successfully ablated microglia and rescued the loss of *Dek*-silenced ECII neurons. We next tested the longer-term effects of *Dek* silencing on somatodendritic tau accumulation in ECII neurons. We compared the somatic levels of phospho-tau (Thr231) in the layer II of the EC between control and *Dek* silenced entorhinal cortices in PLX5622 treated mice and found a significant accumulation of tau after *Dek* silencing (**Figure 6b**), suggesting that DEK is a crucial regulator of tau accumulation in the neurons most vulnerable to AD.

## DISCUSSION

In the present work we set out to identify molecular processes underlying selective neuronal vulnerability in AD by adopting a cell-type specific point of view. We used an *in silico* model of ECII neurons; followed up on our predictions in these same neurons *in vitro* and *in vivo*; and analyzed the effects of gene perturbation in a cell-type specific way *in vivo* using bacTRAP. We demonstrate the strong predictive performance of the *in silico* networks by comparing DEK neighbors in ECII functional networks and genes differentially expressed in these very cells after *Dek* silencing. Previous studies based on functional screens have highlighted very important regulators of tau accumulation *in vitro* (Lasagna-Reeves et al., 2016; van der Kant et al., 2019; Wang et al., 2017). A feature of our framework compared to these studies is that it is the first one to integrate quantitative genetics data (Beecham et al., 2014), in addition to *in vivo* genomics data for vulnerable neurons (Lasagna-Reeves et al., 2016; van der Kant et al., 2019; Wang et al., 2017). It is thus likely to identify genes that are indeed regulating tau during the natural course of AD within ECII neurons, especially during early disease stages. In the present study, by searching for upstream regulators of the AD vulnerability module, we demonstrate that *DEK*, the top gene identified by our data-driven approach, is a regulator of total tau levels *in vitro*, and most importantly *in vivo* in a human tau mouse model. Interestingly, a single nucleotide polymorphism in a *DEK* intron, rs145578678, was recently found to have a suggestive association with the rate of cognitive decline in AD (p-value = 3E-6) (Sherva et al., 2020), indicating that genetic variation in DEK could alter the odds of late-onset AD. Even though *DEK* expression levels have not been found to be altered in neurons in recent single-cell transcriptomics studies in human AD (Mathys et al., 2019), DEK function can be regulated by posttranslational modifications, such as phosphorylation (Kappes, Damoc, et al., 2004) and by degradation (Babaei-Jadidi et al., 2011). Further studies will be necessary to elucidate how alterations in DEK function could contribute to ECII vulnerability, and if any known AD drivers (e.g., amyloid pathology, aging or ApoE4) can lead to these alterations.

While the role of DEK as a chromatin modulator in cancer has been extensively studied, its function in neurons has not been previously described, apart from studies documenting expression in the healthy mouse brain and in patients with schizophrenia (Ghisays et al., 2018; O’Donovan et al., 2018). We find that one of the most striking effects of *Dek* silencing is on IEG expression, reminiscent of links between IEGs (e.g. Npas4 and Egr1) and AD pathogenesis (Bartolotti, Segura, & Lazarov, 2016; Castillo et al., 2017; Hu et al., 2019; Miyashita et al., 2014). Somewhat paradoxically, while under basal conditions *Dek* silencing is accompanied by decreased expression of IEGs, *Dek*-silenced neurons show a higher induction of these same genes by KCl. KCl induces neuronal depolarization in a manner that is independent from synaptic transmission. We thus think that higher KCl-evoked IEG induction might be the primary effect of DEK. Sustained aberrant IEG expression could then be interpreted by transduced neurons as sustained synaptic stimulation. It would trigger homeostatic changes to normalize this activity, like for instance downregulation of all AMPA glutamate receptors which we observe in our cultures (GluR2 (*Gria2*), FDR = 1.68E-52, logFC –1.22, GluR3 (*Gria3*), FDR = 6.91E-09, logFC = –0.57, GluR1 (*Gria1*) FDR = 4.29E-07, logFC = −0.43, GluR4 (*Gria4*) FDR = 1.23E-4, log FC = −0.35). In turn, this downregulation would drastically reduce the stimulation of transduced neurons, and eventually decrease IEGs expression. Our ATAC-seq data showed increased chromatin accessibility at the proximal enhancer and promoter regions of the IEG *Egr1*, suggesting that *Dek* silencing epigenetically primes cells for a larger induction of these genes. This effect was accompanied by differences in the total levels of several histone marks although the lack of specific antibodies hampered their mapping to specific locations in the genome. The prediction that DEK is a major hub of the AD vulnerability and axon plasticity module suggests that it might serve as a broad epigenetic modulator of synaptic homeostasis and remodeling in ECII neurons, with potentially long-lasting effects compared to purely transcriptional regulators. Detrimental effects triggered by DEK dysregulation, like tau accumulation, could unfortunately be durable as well.

Tau accumulation in *Dek*-silenced neurons was not accompanied by its hyperphosphorylation or by changes in its transcription, which suggests that it is mediated by impaired translation or degradation. For instance, tau translation can be modulated by the activation of AMPA and NMDA receptors (Kobayashi, Tanaka, Soeda, Almeida, & Takashima, 2017), and alterations in NPAS4 levels may affect tau degradation (Fan et al., 2016). Also, tau acetylation modulates its turnover (Caballero et al., 2021; Choi et al., 2020; Cook et al., 2014; Min et al., 2015) and affects its aggregation (Carlomagno et al., 2017; Cohen et al., 2011). Indeed, some authors have suggested that tau acetylation could be an early event in AD (Lucke-Wold et al., 2017) that contributes to neurodegeneration (Min et al., 2015). Interestingly, our RNA-seq data showed a highly significant decrease in the expression of several histone deacetylases (HDACs) in *Dek*-silenced neurons, including *Hdac5*, which is both nuclear and cytosolic (padj = 6.28E-23), and might lead to increased tau acetylation in these neurons. Finally, changes in neuronal activity can increase intracellular tau levels due to alterations in its release (Pooler, Phillips, Lau, Noble, & Hanger, 2013; Yamada et al., 2014). Two weeks after *Dek* silencing we do not observe NFT-like structures in ECII neurons. Longer silencing time might have eventually caused the formation of aggregates. Nonetheless, while NFTs are an unquestioned hallmark of AD, several studies have highlighted the contribution of soluble tau accumulation to neurodegeneration. Indeed, tau lowering compounds were shown to prevent cognitive deficits and neuronal death in different mouse models of tauopathy (Lasagna-Reeves et al., 2016; Santacruz et al., 2005) and have been explored as therapeutic strategy (Wang et al., 2017). Moreover, tau accumulation could be at least partially responsible for the increased intrinsic excitability we found by patch-clamp after *Dek* silencing (Holth et al., 2013; Shao et al., 2022; Sohn et al., 2019). Future in-depth analysis of the crosstalk between neuronal activity and tau levels and post-translational modifications in *Dek*-silenced neurons might contribute to identify mechanisms leading to tau pathology in AD.

A last crucial feature of the silencing of DEK on ECII neurons is its indirect effect on microglia. DEK downregulation *in vivo* leads to the progressive loss of ECII neurons and the accumulation of microglia in the layer II of EC, often in very close contact with the *Dek*-silenced neurons. We observed increased expression of *Ccl3*, a cytokine that is upregulated in microglia in response to neuronal activity (Badimon et al., 2020), which suggests that *Dek* silencing-induced alterations in neuronal excitability elicit changes in microglia function. This was accompanied by increased expression of known mediators of neuroinflammation, synaptic spine loss (*Itgam, C1qa)* (Hong et al., 2016; Lui et al., 2016; Merlini et al., 2019; Vasek et al., 2016) and neuronal loss (*Cx3cr1*) (Fuhrmann et al., 2010). Furthermore, we also observed increased levels of *Tyrobp* a driver gene for AD (Zhang et al., 2013) and *Spp1*, both upregulated in the disease-associated microglia that is found in AD (Mathys et al., 2019; Olah et al., 2020). Altogether these results suggest that altered neuronal activity mediated by DEK loss of function triggers neuron-to-microglia signaling mechanisms that steer microglia towards a pro-inflammatory and phagocytic state that contributes to neurodegeneration. Indeed, in agreement with this notion, depletion of microglia with the PLX5622 compound rescued the loss of ECII neurons. This neuron-glia crosstalk regulated by DEK in neurons might take center stage during early AD pathology in the EC.

While uncontestable that reactive microglia and tau accumulation are manifestations of AD, the idea that alterations in network excitability and defects in synaptic homeostasis could also play a role in early disease is gaining traction. Dysregulation of IEGs, like Npas4 and Egr1, has also been linked to AD pathogenesis (Bartolotti, Segura, & Lazarov, 2016; Castillo et al., 2017; Hu et al., 2019; Miyashita et al., 2014) (Frere & Slutsky, 2018; Styr & Slutsky, 2018). Here we uncover DEK as a plausible upstream regulator of all these processes, interconnecting them in a sequence of events that could potentially culminate in the degeneration of ECII neurons during the early stages of AD. Future studies will be essential to fully understand how alterations in DEK function can contribute to neuron vulnerability to AD, and how this novel axis is affected by known drivers of AD.

## ACKNOWLEDGEMENTS

We thank Paul Greengard for his unwavering support of the early stages of this project, until his sudden passing. We thank C. Zhao, C. Lai and the whole Rockefeller University genomics resource center team for all the sequencing. We thank T. Carroll and the Rockefeller University Bioinformatics facility for their help with data analysis. P.R-R. was supported by the European Union’s Horizon 2020 research and innovation program under the Marie Sklodowska-Curiegrant agreement No 799638. P.R-R and C.T. were supported by Alzheimerfonden and Margaretha af Ugglas Stiftelse. P.R-R., M.F. and J.P.R. were supported by the Fisher Center for Alzheimer’s Disease Research. J.P.R. was supported by Cure Alzheimer’s Fund. This study was supported by the National Institute on aging of the NIH (awards RF1 AG054564 and RF1 AG047779 to J.P.R.).

## AUTHORS CONTRIBUTION

Authors contribution is assigned following CRediT (Contributor Roles Taxonomy) guidelines. Conceptualization and methodology, J.P.R and P.R-R.; Formal Analysis, W.W, V.Y and L.L.; Investigation, P.R-R, L.E.A-G, L.L, C.T, V.Y and J.P.R.; Resources, I.S-A, Z.P, S.S, A.C-M, M.F, V.Y and J.P.R.; Writing-Original Draft, J.P.R and P.R-R.; Writing-Review & Editing, L.E.A-G, L.L, A.C-M, O.T, M.F and V.Y.; Visualization, P.R-R, L.E.A-G and L.L.; Funding Acquisition, J.P.R, P.R-R, A.C-M and M.F.

## METHODS

### Mouse models

ECII-bacTRAP mouse were previously generated in our laboratory (Roussarie et al., 2020). They transgenically express the BAC #RP23-307B16 where a cDNA encoding eGFP-L10a was integrated before the start codon of Sh3bgrl2. As a result, they express eGFP-L10a under the control of the regulatory regions of the ECII enriched Sh3bgrl2 gene, which leads to eGFP-L10a expression exclusively in ECII neurons. Human-tau transgenic mice (hMAPT mice) (Andorfer et al., 2003) were obtained from Jackson (B6.Cg-Mapttm1(EGFP)Klt Tg(MAPT)8cPdav/J, Strain #005491). Experiments were performed in 12-15 months of age mice. The groups were age-matched for each experiment.

Mice were maintained on a 12h dark/light cycle and provided with rodent diet and water *ad libitum*. All the experiments involving mice were approved by the Rockefeller University Institutional Animal care and Use Committee (IACUC protocols #16902 and 19067-H).

### Mouse primary neuron cultures

Entorhinal cortex or cortico-hippocampal (for H3.3 and H3K27ac CUT&Tag) neurons in primary culture were prepared from embryos of C57Bl/6J mice (Jackson, stock# 000664). Briefly, entorhinal cortices from E17 mice were incubated at 37°C in 0.05% trypsin/EDTA (#T11493, ThermoFisher) for 10 min. After centrifugation the tissue pellet was dissociated in HBSS containing 0.5mg/ml DNAse I (#10104159001, Merck) with a glass Pasteur pipette. Cells were seeded at 50,000 cells/cm^2^ in Neurobasal medium (#21103049, ThermoFisher), supplemented with 2% B-27 (#17504044, ThermoFisher) and 2 mM GlutaMAX (#35050061, ThermoFisher), and grown at 37 °C in a humidified 5% CO_2_-containing atmosphere. All experiments were performed after a minimum of 10 days *in vitro* (DIV). The experiments involving primary cultures were approved by The Rockefeller University (IACUC protocols #16902 and 19067-H) and Karolinska Institute (4884-2019) ethical committees.

### AAVs

Purified Adeno-associated virus (AAVs) stocks were produced by Vector Biolabs. The viruses used for *Dek* silencing were: AAV1-mCherry-U6-mDEK-shRNA (shRNA sequence: 5’-CCGG-CGAACTCGTGAAGAGATCTTCTCGAGAAGATCCTCTTCACGAGTTCG-TTTTT-3’) and AAV1-mCherry-U6-scrmb-shRNA (control shRNA sequence: 5’- CCGG-CAACAAGATGAAGAGCACCAACTCGAGTTGGTGCTCTTCATCTTGTTG-TTTTT-3’). The viruses used for *Dek* overexpression were: AAV1-hSyn1-mDEK-IRES-mCherry (expressing the cDNA encoding for mouse DEK – NM_025900) and AAV1-hSyn1-mCherry-WPRE (empty control).

### EC mouse primary neuron treatments

#### AAVs transduction

EC neurons in primary culture at 7 days *in vitro* (DIV) were transduced by a full media change to media containing either control, mDEK-IRES or mDEK-shRNA AAVs at a concentration of 1 × 10^10^ GC/ml of media. Neurons were maintained in virus-containing media until the day of the experiment.

#### KCl stimulation

EC neurons were stimulated by a full media change to media containing either vehicle (PBS) or 50 mM KCl. Neurons were incubated with KCl for the whole duration of the experiment.

### ATAC-seq

ATAC-seq was performed as previously described (Buenrostro, Wu, Chang, & Greenleaf, 2015). Briefly 50,000 EC primary neurons were pelleted by centrifugation (5 min, 500 xg at 4°C) and lysed in 10 mM Tris-HCl pH 7.4, 10 mM NaCl, 3 mM MgCl_2_, 0.1% NP-40, 0.1% Tween-20 and 0.01% digitonin. Nuclei where then pelleted by centrifugation (10 min, 1000 xg at 4°C) and resuspended in transposition reaction mix: 1% digitonin, 10% Tween-20, Tn5 transposase and TD buffer (Nextera DNA Library Prep Kit, #FC-121-1030, Illumina). After transposase reaction (30 min, 37°C, 1000 RPM) and DNA purification (MinElute Reaction Cleanup Kit, #28204, Qiagen), the transposed samples were amplified by PCR with individual barcode primers per sample (NEB Next High-Fidelity 2x PCR Master Mix, #M0541S, NEB). After 5 PCR cycles, a fraction of the partially amplified library was quantified by qPCR to calculate the number of additional PCR cycles needed to amplify each sample without saturation (number of cycles needed to reach 1/3 of the maximum R). The amplified libraries were purified with Agencourt AMPure XP magnetic beads (#A63880, Beckman Coulter) and their quality and concentration were determined with an Agilent High Sensitivity DNA bioanalysis Chip. Libraries were sequenced on a NextSeq sequencer (Illumina).

### Mass spectrometry analysis of histone posttranslational modification

A mass spectrometry analysis of histone posttranslational modification (Mod-spec) was performed by Activ Motif. Bulk histones were acid-extracted from pellets obtained from EC neurons in primary culture transduced with either control or *Dek*-silencing AAVs, propionylated and subjected to trypsin digestion as described previously (Zheng, Thomas, & Kelleher, 2013). Briefly, histones were extracted at room temperature for 1 hour in 0.2 M sulfuric acid with intermittent vortexing. Histones were then precipitated by the addition of trichloroacetic acid (TCA) on ice, and recovered by centrifugation (10,000 x g, 5 min at 4°C). The pellet was then washed once with 1mL cold acetone/0.1% HCl, twice with 100% acetone, and air dried. Histones were propionylated with 1:3 v/v propionic anhydride/2-propanol and incremental addition of ammonium hydroxide to keep the pH around 8, and subsequently dried in a SpeedVac concentrator. The pellet was then resuspended in 100 mM ammonium bicarbonate and adjusted to pH 7-8 with ammonium hydroxide. Histones were then digested with trypsin resuspended in 100 mM ammonium bicarbonate overnight at 37°C and dried in a SpeedVac concentrator. The pellet was resuspended in 100 mM ammonium bicarbonate and propionylated a second time as described above. Histone peptides were resuspended in 0.1% TFA in H_2_O for mass spectrometry analysis.

Samples were analyzed on a triple quadrupole (QqQ) mass spectrometer (Thermo Fisher Scientific TSQ Quantiva) directly coupled with an UltiMate 3000 Dionex nano-liquid chromatography system. Peptides were first loaded onto an in-house packed trapping column (3cm×150μm) and then separated on a New Objectives PicoChip analytical column (10 cm×75 μm). Both columns were packed with New Objectives ProntoSIL C18-AQ, 3μm, 200Å resin. The chromatography gradient was achieved by increasing percentage of buffer B from 0 to 35% at a flow rate of 0.30 μl/min over 45 minutes. Solvent A: 0.1% formic acid in water, and B: 0.1% formic acid in 95% acetonitrile. The QqQ settings were as follows: collision gas pressure of 1.5 mTorr; Q1 peak width of 0.7 (FWHM); cycle time of 2 s; skimmer offset of 10 V; electrospray voltage of 2.5 kV. Targeted analysis of unmodified and various modified histone peptides was performed. This entire process was repeated three separate times for each sample.

#### Data analysis

Raw MS files were imported and analyzed in Skyline with Savitzky-Golay smoothing (MacLean et al., 2010). All Skyline peak area assignments for monitored peptide transitions were manually confirmed. Multiple peptide transitions were quantified for each modification. For each monitored amino acid residue, each modified (and un-modified) form was quantified by first calculating the sum of peak areas of corresponding peptide transitions; the sum of all modified forms was then calculated for each amino acid to represent the total pool of modifications for that residue. Finally, each modification is then represented as a percentage of the total pool of modifications. This process was carried out for each of the three separate mass spec runs, and the raw data provided in the data delivery spreadsheet corresponds to the mean and standard deviation of the resulting three values from this analysis for each modified and unmodified form of the corresponding amino acid residue.

### CUT&Tag

Frozen cell pellets from control and *Dek*-silenced EC (For H3K36Ac) or cortico-hippocampal (For H3.3 and H3K27Ac) mouse primary neurons were sent to Active Motif for CUT&Tag. Briefly, cells were washed and incubated over-night with Concanavalin A beads and 1.3 µl/reaction of primary antibodies against H3K27Ac (#39133, Active Motif), H3.3 (#91191, Active Motif) and H3K36Ac (#39379, Activ Motif). After incubation with the secondary anti-rabbit antibody (1:100), cells were washed and tagmentation was performed at 37°C using protein-A-Tn5. Tagmentation was halted by the addition of EDTA, SDS and proteinase K after which DNA extraction and ethanol purification was performed, followed by PCR amplification and barcoding (see Active Motif CUT&Tag kit, catalog number 53160 for recommended conditions and indexes). Following SPRI bead cleanup (Beckman Coulter), the resulting DNA libraries were quantified and sequenced on Illumina’s NextSeq 550 (8 million reads, 38 paired end).

#### Data analysis

The paired-end 38 bp sequencing reads generated by Illumina sequencing were mapped to the genome using the BWA algorithm with default settings. Only reads that passed Illumina’s purity filter, aligned with no more than 2 mismatches, and mapped uniquely to the genome were used to subsequent analysis. Duplicate reads were removed. Genomic regions with high levels of transposition/tagging events were determined using the MACS2 peak calling algorithm. To identify the density of transposition events along the genome, the genome was divided into 32 bp bins and the number of fragments in each bin was determined by extending the reads to 200 bp, which is close to the average length of the sequenced library inserts. To compare peak metrics between samples, overlapping intervals were grouped into “merged regions” which are defined by the start coordinate of the most upstream interval and the end coordinate of the most downstream interval. In locations where only one samples has an interval, that interval defined the merged region.

### Electrophysiology

Mouse EC neurons in primary culture were grown on glass coverslips. Coverslips were placed in a submerged chamber perfused with aerated artificial cerebrospinal fluid: 124 mM NaCl, 30 mM NaHCO_3_, 10 mM glucose, 1.25 mM NaH_2_PO_4_, 3.5 mM KCl, 1.5 mM MgCl_2_, 1.5 mM CaCl_2_, at 36°C with a perfusion rate of 1-2 ml per minute. Coverslips were left undisturbed for 5 minutes before any recording. Patch-clamp (whole-cell) recordings were performed with borosilicate glass microelectrodes (4–6 MΩ) from the soma of visually identified pyramidal cells using IR-DIC microscopy (Zeiss Axioskop, Germany) using a Multiclamp 700B (Molecular Devices, CA, USA). For neuronal electrophysiological properties and spontaneous excitatory postsynaptic current (sEPSC) measurements (Vh = −70 mV) a potassium-based intracellular solution was used: 122.5 mM K-gluconate, 8 mM KCl, 4 mM Na_2_-ATP, 0.3 mM Na_2_-GTP, 10 mM HEPES, 0.2 mM EGTA, 2 mM MgCl_2_, 10 mM Na_2_- Phosphocreatine, set to pH 7.2-7.3 with KOH, osmolarity 270-280 mOsm. The signals were sampled, and low pass filtered at 2 kHz, digitized, and stored using a Digidata 1440A and pCLAMP 10.4 software (Molecular Devices, CA, USA).

#### Resting membrane potential

(RMP) was determined in current-clamp mode after breaking the membrane and calculate from the mean of 1 min recording using Clampfit11.2. Cell capacitance (Cm) and Tau (τ) values were taken directly from the Clampex membrane test tool and amplifier readings. Input resistance was measured from subthreshold hyper- and depolarizing current steps (−20 to 20 pA, 10 pA increments, 700 ms; Vh = −70 mV) and calculated using Clampfit11.2. Rheobase: First-spike latency measurements were done by applying a rheobase protocol to PCs in current-clamp whole cell configuration to elicit the firing of a single AP (10 pA increments, 300 ms). The current at which the first AP was fired was define as current threshold. First-spike latency was then calculated as the time between the beginning of the test current pulse and the maximus amplitude of the AP. Sag: A hyperpolarizing current pulse was applied to PCs to investigate the SAG (Vh = −60 mV). Amplitude was calculated from the steady state of the current to the peak. AP waveform: Long-lasting current steps to evoke 1 AP were used to analyze AP properties (10 pA increments, 700 ms). AP parameters (amplitude, firing threshold, half-width, maximum rise slope (MRS), time to MRS, maximum decay slope (MDS) and time to MDS), AP waveform plot and analysis were performed in Clampfit11.2. sEPSC were detected off-line using MiniAnalysis software (Synaptosoft, Decatur, GA, USA). Charge transfer, event amplitude and event frequency were analyzed using Microsoft Excel (Microsoft Office) and GraphPad Prism (GraphPad Software, USA) with the result representing average values taken over 1 min periods.

### Stereotaxic injections

AAVs were injected in the entorhinal cortex of adult mice using an Angle Two mouse stereotaxic frame with a motorized nanoinjector (Leica). Animals were anesthetized with xylazine (4.5 mg/kg body weight) and ketamine (90 mg/kg body weight) injected peritoneally. An ophthalmic ointment was applied to the anesthetized animals to prevent corneal drying during the procedure. AAVs were loaded in a 10 ml syringe (#7653-01, Hamilton) with a 33-gauge needle (#7803-05, Hamilton). 2 ml of AAVs (1 × 10^13^ GC/ml) were injected in the mouse EC (AP: −3.70; ML: −4.65; DV: −4.50) with the nanoinjector tilted −4.05°. The same volume and concentration of control AAVs were injected in the contra-lateral EC (AP: −3.70; ML: +4.65; DV: −4.50) with a nanoinjector tilt of +4.05°. Injection rate was 0.4 ml/min. Bacitracin antibiotic gel was applied to the surgery wound, that was sutured with a non-absorbable monofilament. To compensate fluid loss, warm sterile saline solution was injected intraperitoneally (3% of the body weight), and animals were kept on a heated pad and monitored until complete recover from anesthesia.

### RNA-seq

#### Primary culture RNAseq

For RNAseq analysis of EC primary cultures, RNA was extracted 4 days after AAV transduction using the Purelink RNA kit. 200ng of RNA was used to generate RNAseq libraries using the Truseq RNA sample prep kit. Libraries were sequenced using a NextSeq 500 Illumina sequencer.

#### bacTRAP-RNAseq

Entorhinal cortices from both hemispheres were isolated from individual mice and processed for bacTRAP as previously described (Heiman et al., 2014; Roussarie et al., 2020). Briefly, tissue was homogenized on 1ml of lysis buffer (20 mM Hepes KOH, 10 mM MgCl_2_, 150 mM KCl, 0.5 mM DTT, 100 ug/ml cycloheximide) supplemented with protease (#A32965, ThermoFisher) and RNAse inhibitors (40U/ml RNAsin, #N2515, Promega and 20U/ml Superasin, #AM2696, ThermoFisher) at 4°C in a glass Teflon homogenizer. After centrifugation, the supernatant was incubated with 1% NP-40 and 30 mM DHPC (#850306P, Avanti) on ice for 5 minutes. After centrifugation, the supernatant, containing ribosome bound RNAs, was incubated over-night with magnetic beads (Streptavidin MyOne T1 Dynabeads, #65602, ThermoFisher) previously coated with anti EGFP antibodies for immunoprecipitation (HtzGFP-19C8 and HtzGFP-19F7, from MSKCC monoclonal antibody facility (Heiman et al., 2008)). Immunoprecipitated RNAs were then purified using the RNeasy Plus Micro Kit (#74034, Qiagen). RNA integrity was determined with a Bioanalyzer 2100 (Agilent) using an RNA 6000 pico chip (#5067-1513, Agilent). RNA was quantified with Quant-it Ribogreen RNA reagent (#R11490, ThermoFisher). Reverse transcription was performed with Ovation RNAseq v2 kit (#7102, NuGEN) from 5 ng of RNA following the manufacturer’s instructions. cDNAs were purified using the QIAquick PCR purification kit (#28104, Qiagen). cDNA yield was measured with Quant-IT Picogreen dsDNA kit (#P7581, ThermoFisher). 200 ng of cDNA were used for fragmentation prior to cDNA library preparation. cDNA was sonicated into 200 bp fragments using a Covaris S2 ultrasonicator instrument (10% duty cycle, intensity 5, 200 cycles/burst per second for 2 minutes at 5.5° to 6°C). Library preparation was performed with the TruSeq RNA sample preparation kit v2 (#RS-122-2001, Illumina) and were sequenced at the Rockefeller University genomics resource center on a NextSeq 500 sequencer (Illumina).

### PLX5622 treatments

PLX5622 compound was obtained from MedChemExpress (New Jersey, USA). Chow containing PLX5622 at 1200 ppm was manufactured by a trained diet preparation operator at Research Diets. Inc (New Jersey, USA) in AIN-76A rodent diet. Animals were supplied with AIN-76A diet pellets either alone (control) or containing PLX5622 *ad libitum*. Diet supplementation was initiated 5 days before stereotaxic injections were performed and was maintained for the whole duration of the experiments.

### Mouse tissue processing

To allow for dual analysis of mouse tissue samples by western blot and RT-qPCR, dissected entorhinal cortices were homogenized in 100 ml H_2_O supplemented with protease (#A32965, ThermoFisher), phosphatase (#4906837001, PhoSTOP, Merck) and RNAse (40U/ml RNAsin, #N2515, Promega and 20U/ml Superasin, #AM2696, ThermoFisher) inhibitors with a cordless motor pestle homogenizer. Afterwards 45 ml of the lysate was transferred to an RNAse-free tube, mixed with RNA extraction buffer and cleaned up using the Purelink RNA kit (see below). The remaining volume was lysed in RIPA protein extraction buffer (#89901, ThermoFisher).

For immunofluorescence analysis, mice were anesthetized with nembutal and perfused with PBS followed by 4% paraformaldehyde (PFA). Brains were then dissected and post-fixed in 4% PFA for 1h. Afterwards they were washed with PBS and cryopreserved by incubation in increasing sucrose concentrations (5%, 15% and 30%). They were then embedded in OCT compound (TissueTek), cut in 40 mm-thick free-floating horizontal sections with a CM3050 Cryostat (Leica) and further preserved in cryoprotectant solution (50% ethylene glycol, 20% glycerol in PBS) at −20°C until further analysis.

### Western blot

Cells or mouse brain tissue was homogenized in RIPA buffer (#89901, ThermoFisher), supplemented with protease (#A32965, ThermoFisher), and phosphatase (#4906837001, PhoSTOP, Merck) inhibitors. The cell extracts were then subjected to sodium dodecyl sulfate (SDS) polyacrylamide gel electrophoresis and transferred to a nitrocellulose membrane. Membranes were blocked in Intercept (TBS) blocking buffer (#927-60001, Li-Cor) 1h at room temperature and blotted with the primary antibodies overnight at 4 °C. Antibodies used were AT8 (1:2000, #MN1020, ThermoFisher), Tau (1:1000, #A0024, Dako) and Iba1 (1:1000, #019-19741, Wako). Beta actin (1:1000, #4967, Cell Signaling) was used as loading control. The day after membranes were incubated with IRDye 800CW and 680RD mouse and rabbit secondary antibodies (Li-cor) for 1h at room temperature. Signal detection was performed with a Oddisey scanning system (Li-Cor). Band intensity quantification was performed with the Image Studio Lite software.

### Immunofluorescence

#### EC neurons in primary culture

were seeded in Poly-D-ly-sine-coated round glass coverslips. After the treatments, they were fixed in 4% PFA for 10 minutes. Fixed cells were then blocked for 30 min in 1% bovine serum albumin (BSA), 0.1% Triton X-100, phosphate buffered saline (PBS, Thermo-Fisher, Massachusetts, USA) and incubated with the primary antibody overnight at 4ºC. The antibodies used were AT8 (1:1000, #MN1020, ThermoFisher) and MAP2 (1:1000, #ab183830, Abcam). The day after, the cells were washed with PBS and incubated with the secondary antibody in PBS for 1h at room temperature. The secondary antibodies used were Goat anti-mouse IgG (H+L) Cross-adsorbed Alexa fluor 488 (#A-11001, ThermoFisher) and Goat anti-rabbit IgG (H+L) Cross-adsorbed Alexa fluor 594 (#A-11012, ThermoFisher). DAPI was used as nuclear staining. Cells were then washed and mounted with ProLong Gold Antifade Reagent (#P36980, ThermoFisher).

### 40 mm-thick free-floating horizontal sections

were permeabilized in 0.1% fish gelatin, 2% normal goat serum and 0.1% Triton X-100 in PBS for 30 min at room temperature. They were then stained over night at 4°C with primary antibodies diluted in the same permeabilization buffer. Primary antibodies used were mCherry (1:1000, #ab205402, Abcam), Iba1 (1:1000, #019-19741, Wako), Reelin (1:500, #MAB5364, Millipore), AT180 (1:1000, #NB100-82249, Novus Bio). The day after samples were washed in PBS and incubated with secondary antibodies for 1h at room temperature. The secondary antibodies used were Goat anti-mouse IgG (H+L) Cross-adsorbed Alexa fluor 488 (#A-11001, ThermoFisher), Goat anti-chicken IgG (H+L) Cross-adsorbed Alexa fluor 594 (#A32759, ThermoFisher) and Goat anti-rabbit IgG (H+L) Cross-adsorbed Alexa fluor 647 (#A-21245, ThermoFisher). After washing with PBS auto-fluorescence was quenched with Eliminator reagent (#2160, Millipore Sigma) following the manufacturer’s instructions and mounted with ProLong Gold Antifade Reagent (#P36980, ThermoFisher).

### Image acquisition

All images were acquired on a Zeiss LSM 510 META laser scanning confocal microscope.

### For Iba1 and AT180 fluorescence intensity quantification

in mouse sections, a minimum of 3 pictures were taken from different sections and from each hemisphere from the same mouse (shDEK and Control transduced). The region of interest was established manually by selecting the entorhinal cortex layer II and on slides where mCherry positive cells (AAV transduced) could be detected. Iba1 and At180 fluorescence intensity was determined using the threshold selector to identify positive staining signal.

### RT-qPCR

#### EC neurons in primary culture

RNA extraction, purification, reverse-transcription, and qPCR were performed using the TaqMan Fast Advanced Cells-to-CT kit (#A35377, ThermoFisher), according to the manufacturer’s instructions.

#### Mouse tissue homogenates

RNA was purified using the PureLink RNA mini kit with in-column DNAse-I digest (#79254, Qiagen) following the manufacturer’s instructions (#12183018A, ThermoFisher). RNA was then quantified in a nanodrop and around 400 ng of RNA was reverse-transcribed using the SuperScript III First-Strand Synthesis System with a 1:1 mix of oligo dT and random hexamers oligonucleotides (#18080051, ThermoFisher), followed by RNAse H digestion. cDNA was then used to run qPCRs using TaqMan Universal PCR master mix.

FAM-labeled TaqMan assays (ThermoFisher) used were: Mm00662582_m1 (*Dek*), Mm00656724_m1 (*Egr1*), Mm00463644_m1 (*Npas4*), Mm00521992_m1 (*Mapt*), Mm00438354_m1 (*Cx3cr1*), Mm07295529_m1 (*C1qa*), Mm00434455_m1 (*Itgam*), Mm99999057_m1 (*Ccl3*), Mm00449152_m1 (*Tyrobp*), Mm00444701_m1 (*Ccl24*), Mm00446026_m1 (*P2ry12*) and Mm00436767_m1 (*Spp1*). *Gapdh* was used as endogenous control (4352932E). All qPCRs were performed in a QuantStudio 12K-flex machine.

### Network connectivity analysis

To identify genes that are strongly connected to the AD vulnerability module, for each gene *g* in the entorhinal cortex functional network, we calculated a z-score for vulnerability module connectivity:

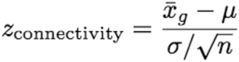

where 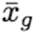 is the mean of posterior probabilities between *g* and all genes in the AD vulnerability module, and *n* is the number of genes in the module. m and s are respectively the mean and standard deviation of posterior probabilities of all genes in the network to gene *g*. Thus, high *z*_*connectivity*_ represents stronger connectivity between gene *g* to the AD vulnerability module than expected based on gene *g*’s general connectivity patterns within the entorhinal cortex network.

### Differential gene expression analysis

Following sequencing, adapter and low-quality bases were trimmed by fastp (S. Chen, Zhou, Chen, & Gu, 2018) from the raw sequencing files in FASTQ format. Cleaned reads were aligned to the mm10 reference genome using STAR version 2.7.1a (Dobin et al., 2013). After alignment, the Reads Per Kilobase of transcript per Million mapped reads (RPKM) for all genes in each sample were calculated with R package edgeR (McCarthy, Chen, & Smyth, 2012). To analyze differential gene expression between samples, DESeq2 (Love, Huber, & Anders, 2014) was used, applying the standard comparison mode between two experimental groups.

