## Supplementary material for "The proto-oncogene DEK regulates neuronal excitability and tau accumulation in Alzheimer’s disease vulnerable neurons": Suplementary Figures and Table S3

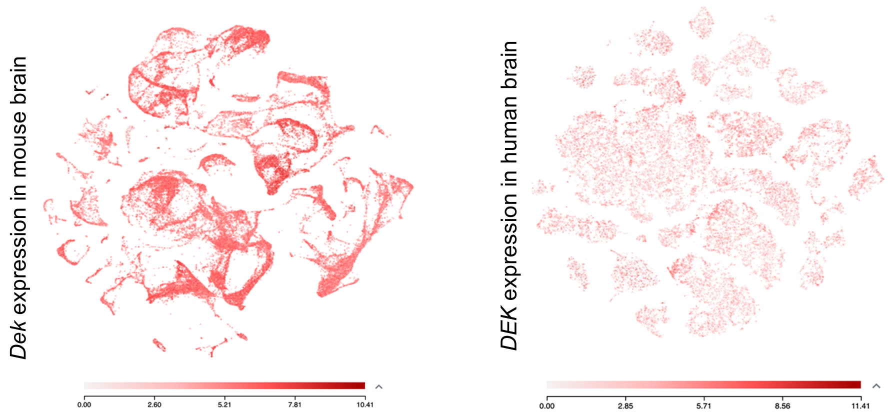

**Figure S1. DEK expression in the mouse and the human brain.** Scatter plots of DEK expression levels in different cell types of the mouse (left panel) and human brain (right panel). Source: Allen brain map transcriptomics explorer.

**A**

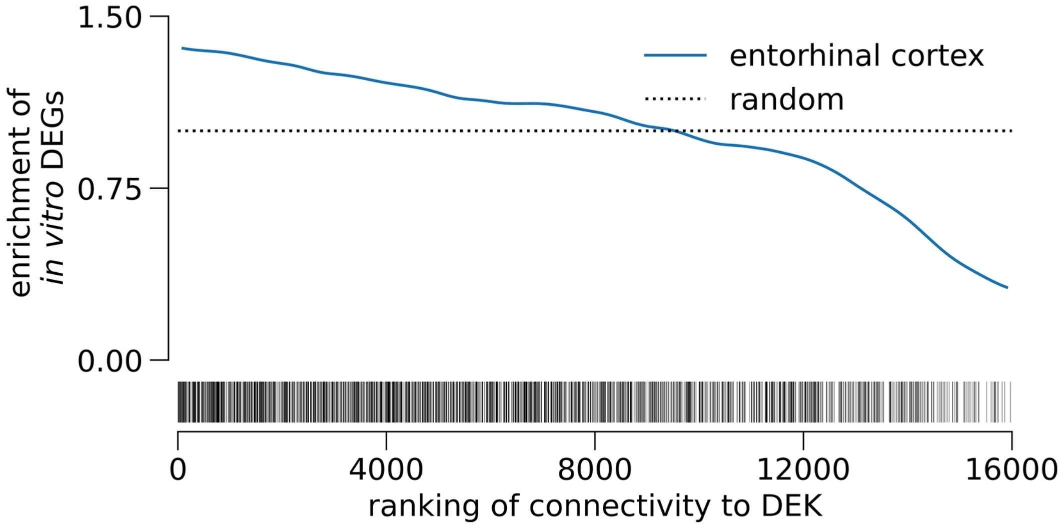

Figure S2. Significant enrichment of genes changing *in vitro* from *Dek*-silencing in *in silico* ECII network predictions. Enrichment analysis of DEGs (FDR<0.05, denoted in the rug plot, bottom) between control and *Dek*-silenced ECII neurons *in vitro* demonstrates strong enrichment within genes ranked by probability of functional interaction with DEK in the entorhinal cortex network (blue). Dashed line represents expected enrichment if given random predictions.

A

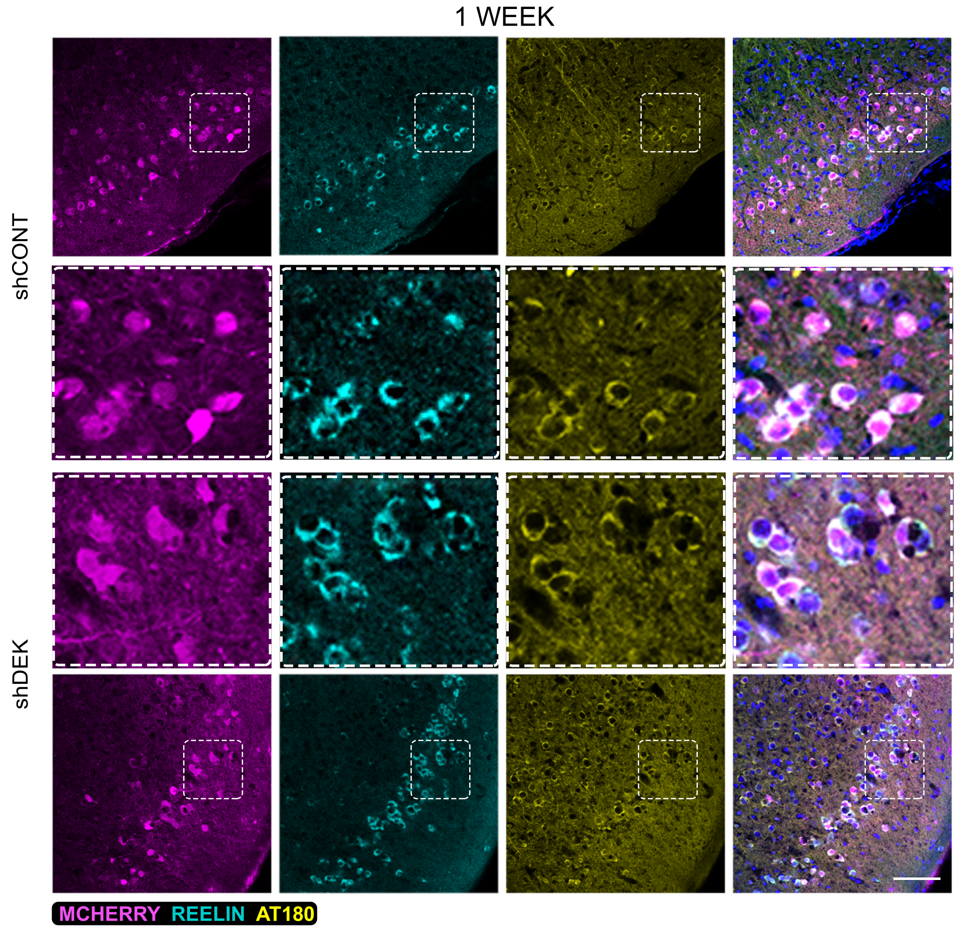

B

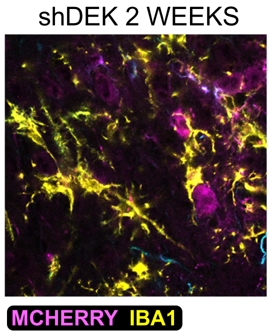

**Figure S3. Effect of Dek silencing in ECII neurons of hMAPT mouse. A)** Confocal microscopy images of immunofluorescence staining of transduced neurons (mCherry, magenta), ECII neurons (Reelin, cyan) and phospho-tau Thr231 (AT180, yellow) at 1-week post-transduction of control or shDek-carrying AAVs in hMAPT mice. Scale bar 100 um**. B)** Confocal microscopy images of immunofluorescence staining of shDek AAVs-transduced neurons (mCherry, magenta) and microglia (Iba1, yellow) at 2 weeks post-transduction

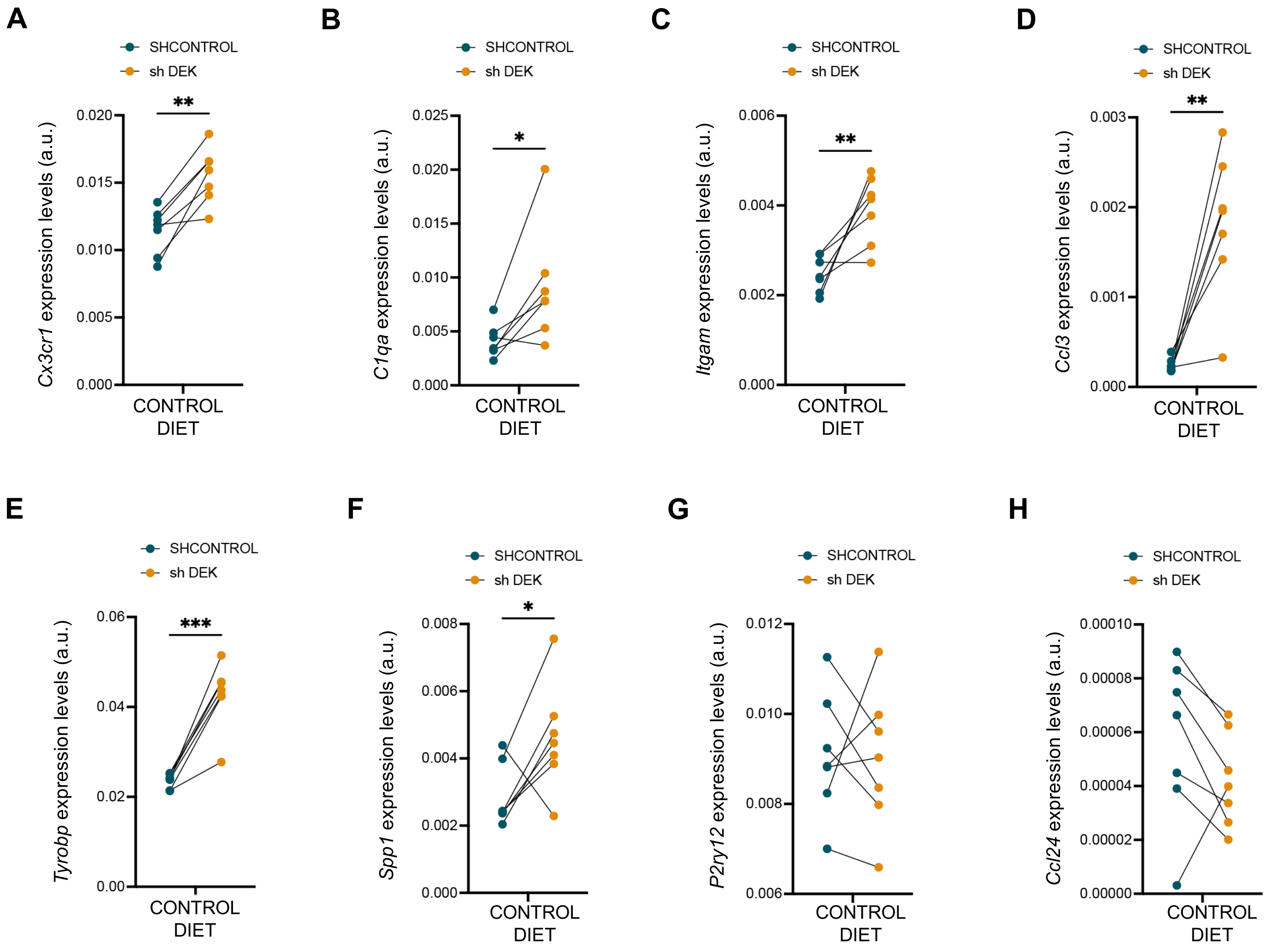

**Figure S4. Effect of Dek-silencing in ECII neurons on microglia gene expression in hMAPT mouse. A-H)** RT-qPCR quantification of the expression levels of *Cx3cr1* **(A)**, *C1qa* **(B)**, *Itgam* **(C)**, *Cd3* **(D)**, *Tyrobp* **(E)**, *Spp1* **(F)**, *P2ry12* **(G)** and *Ccl24* **(H)** in bulk homogenates of the control and the *Dek*-silenced opposite hemisphere for each hMAPT mouse fed with control diet. Paired t-test * pval<0.05, **pval<0.01, ***pval<0.005.

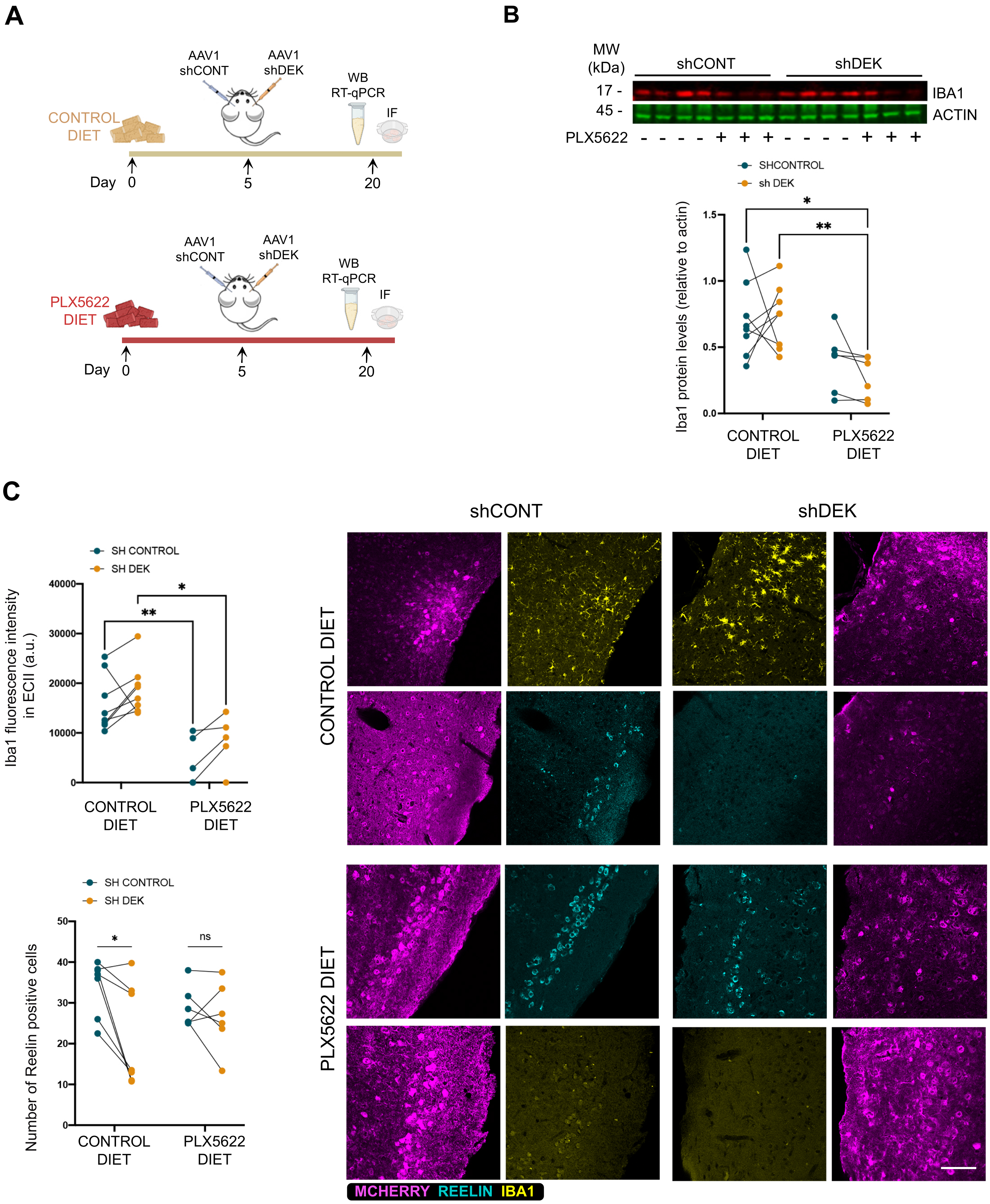

**Figure S5. Effect of Dek silencing in ECII neurons of control and PLX5622 diet treated hMAPT mouse.** **A)** Schematic representation of the experimental layout. **B)** Western blot analysis of Iba1 levels (red) in control or PLX5622 diet fed mice and, in the control, or the Dek-silenced hemisphere of the same representative mice. The graph shows the quantification for the complete mouse cohort, paired by the control and the *Dek*-silenced opposite hemisphere for each mouse. Iba1 levels were normalized by actin. 2way ANOVA Sidak's multiple comparisons test **pval<0.01, *pval<0.05. **C)** Confocal microscopy images of immunofluorescence staining of transduced neurons (mCherry, magenta), ECII neurons (Reelin, cyan) and microglia (Iba1, yellow) at 2 weeks post-transduction of control or PLX5622 diet fed hMAPT mice. Scale bar 100 um. The graphs show the quantification of Iba1 fluorescence intensity (upper panel) and the number of reelin positive cells (lower panel), paired by the control and the *Dek*-silenced opposite hemisphere for each hMAPT mouse. 2way ANOVA Sidak's multiple comparisons test ***pval<0.005, *pval<0.05.

| Electrophysiological properties | Control (n=10) | shDEK (n=11) | p value |
|---|---|---|---|
| **Passive properties** | | | |
| Resting Membrane Potential (mV) | -66.97 ± 6.001 | -59.05 ± 8.321 | **0.0227** |
| Membrane Capacitance (pF) | 17.56 ± 3.531 | 16.26 ± 4.573 | 0.4786 |
| τ (μs) | 624.8 ± 124.2 | 682.6 ± 298.5 | 0.5944 |
| Input Resistance (MΩ) | 159.4 ± 59.17 | 115.1 ± 24.19 | **0.0336** |
| Sag Amplitude (mV) | 2.595 ± 1.747 | 9.407 ± 9.321 | **0.0495** |
| **Active properties** | | | |
| Rheobase | | | |
| Current threshold (pA) | 60.00 ± 24.13 | 44.55 ± 22.96 | **0.0368** |
| First-spike Latency (ms) | 48.83 ± 17.35 | 75.53 ± 40.45 | 0.0824 |
| AP waveform | | | |
| Amplitude (mV) | 50.38 ± 6.823 | 54.19 ± 9.196 | 0.1621 |
| Firing threshold (mV) | -31.20 ± 2.513 | -26.34 ± 4.144 | **0.0046** |
| Half-width (ms) | 2.662 ± 0.4964 | 3.158 ± 0.7493 | 0.0930 |
| Max Rise Slope (mV/ms) | 52.46 ± 23.79 | 35.81 ± 16.75 | 0.0771 |
| Time to Max Rise Slope (ms) | 4.880 ± 0.3743 | 5.150 ± 0.2 | 0.0503 |
| Max Decay Slope (mV/ms) | -21.91 ± 6.245 | -16.79 ± 6.214 | 0.0753 |
| Time to Max Decay Slope (ms) | 7.320 ± 0.4523 | 7.750 ± 0.718 | 0.1265 |
| **Spontaneous Excitatory Input** | | | |
| sEPSC amplitude (pA) | 26.27 ± 13.44 | 17.44 ± 6.515 | 0.0668 |
| sEPSC Frequency (Hz) | 2.667 ± 1.706 | 2.070 ± 0.9253 | 0.3251 |
| sEPSC Charge Transfer (pC) | 17.97 ± 12.79 | 9.465 ± 6.833 | 0.1971 |

**Table S3.** Electrophysiological properties describing active, passive, and excitatory input in control and *Dek*-silenced neurons. Bold values indicate statistical significance.
